# PyBrOpS: a Python package for breeding program simulation and optimization for multi-objective breeding

**DOI:** 10.1101/2023.02.10.528043

**Authors:** Robert Z. Shrote, Addie M. Thompson

## Abstract

Plant breeding is a complex endeavor that is almost always multi-objective in nature. In recent years, stochastic breeding simulations have been used by breeders to assess the merits of alternative breeding strategies and assist in decision making. In addition to simulations, visualization of a Pareto frontier for multiple competing breeding objectives can assist breeders in decision making. This paper introduces Python Breeding Optimizer and Simulator (PyBrOpS), a Python package capable of performing multi-objective optimization of breeding objectives and stochastic simulations of breeding pipelines. PyBrOpS is unique among other simulation platforms in that it can perform multi-objective optimizations and incorporate these results into breeding simulations. PyBrOpS is built to be highly modular and has a script-based philosophy, making it highly extensible and customizable. In this paper, we describe some of the main features of PyBrOpS and demonstrate its ability to map Pareto frontiers for breeding possibilities and perform multi-objective selection in a simulated breeding pipeline.

## 1. Introduction

The advent of modern computer processors has made many computationally intensive breeding-related tasks possible in recent years, including the use of simulation. There are four broad use-cases for computer simulation in breeding: evaluating breeding pipelines, assessing the efficacy of new genetic analysis techniques, studying gene networks and genotype by environment interaction, and modeling crop growth and development (Li et al., 2012). Among these four use-cases, breeding pipeline simulation is perhaps the most practical application in a breeding program. Breeding pipeline simulations allow for breeders to estimate genetic gain and diversity metrics for a proposed breeding strategy without having to spend time and resources empirically testing them. By repeating simulations, breeders may compare the expected performance of multiple proposed breeding strategies, aiding in decision-making processes.

In most circumstances, breeding is multi-objective in nature. In plant breeding, characteristics such as yield, standability, seed and fruit composition, resistance to biotic and abiotic stresses, biomass quality, adaptation to mechanization, and photoperiod response are potential breeding objectives (Fehr, 1991). Additionally, maintaining genetic diversity for long-term genetic gain is important (Gorjanc et al., 2018; Akdemir et al., 2019; Allier et al., 2019a). The path to meeting multiple breeding objectives is not straightforward, unfortunately. Historically, tandem selection, independent culling, and index selection methods have been proposed as strategies to meet multiple breeding goals (Bernardo, 2020). Among these three historical selection strategies, index selection methods are generally regarded as being more effective (Bernardo, 2020).

In more recent years, multi-objective numerical optimization and simulation techniques have been proposed as improvements to traditional methods (Akdemir and Sánchez, 2016; De Beukelaer et al., 2017; Akdemir et al., 2019; Allier et al., 2019a; Moeinizade et al., 2020; Amini et al., 2022). For example, Akdemir and Sánchez (2016) combined an inbreeding objective with a mating risk objective related to the mean and variance of progenies using a weighted sum. De Beukelaer et al. (2017) combined a genetic gain objective with a rare allele management objective using a weighted sum. Akdemir et al. (2019) proposed a non-dominated selection strategy and multi-objective versions of optimal contribution selection (Meuwissen, 1997) and genomic mating (Akdemir and Sánchez, 2016). Allier et al. (2019a) combined a cross usefulness objective with an inbreeding objective cognizant of biased parental genome contributions using the ε-constraint method. Moeinizade et al. (2020) proposed a simulation-based look-ahead-selection strategy for multiple traits. Finally, Amini et al. (2022) proposed a multi-trait, L-shaped index selection strategy to address the shortcomings of traditional index selection techniques. Among these cited selection strategies, most transform multiple objectives into a single objective using weight or constraint parameters determined *a priori* and are consequently solved as single-objective optimization problems. In Akdemir and Sánchez (2016) and Akdemir et al. (2019), the authors utilized a truly multi-objective optimization strategy. They performed multiple optimizations over a range of objective weights and identified efficient frontiers for their objectives.

Multi-objective optimization techniques offer two main benefits over traditional selection techniques. First, multi-objective optimization techniques eliminate or reduce the difficulty of assigning weights to breeding objectives by identifying a set of Pareto optimal or near-Pareto optimal, non-dominated solutions from which to choose. A solution is Pareto optimal if there exists no other solution in the selection search space that can improve upon one objective without making another objective worse. A solution is non-dominated in the context of a set of solutions if there exists no other solution in the set that can improve upon one objective without making another objective worse. Pareto optimal solutions and non-dominated solutions are different from each other in that all Pareto optimal solutions are globally optimal and, by definition, must be non-dominated, while a non-dominated solution may only be locally optimal. This is an important distinction in multi-objective optimization since for most real-world applications it is not known whether a solution is globally optimal. With multi-objective optimization, breeders may visualize identified solutions, examine the tradeoffs between various objectives, and choose from the available points without having to assign weights as one would in an index selection. If desired, pseudo-weights, which are normalized to the range [0,1] and indicate the relative importance of an objective among solutions lying along the Pareto frontier, may also be assigned to identified solutions and serve as a more convenient way to make selection decisions (Deb, 2001). Second, multi-objective optimization techniques allow for the identification of Pareto-optimal solutions that do not lie on the convex hull of the selection search space. A search space is considered convex if a line connecting any two points within the search space lies entirely within the search space. A non-convex search space is one where this is not true for all points. As Amini et al., 2022 emphasized, in discrete selection problems where the Pareto frontier is non-convex, it is mathematically impossible to identify Pareto optimal solutions that do not lie along the convex hull using a linear selection index. This is problematic, as there may exist superb solutions in non-convex regions of the Pareto frontier. A good multi-objective optimization algorithm will be able to identify multiple solutions lying in non-convex Pareto frontier regions from which to choose.

It is common to use evolutionary algorithms to solve multi-objective optimization problems. Evolutionary algorithms are particularly suitable to solving multi-objective optimization problems because they are population based, which allows them to find multiple Pareto optimal or near-Pareto optimal solutions in a single run (Coello Coello et al., 2007). Additionally, evolutionary algorithms are less susceptible to getting trapped in local optima and make few assumptions about the shape and continuity of the Pareto frontier (Coello Coello et al., 2007). In contrast, classical multi-objective optimization techniques such as the weighted sum method, the ε-constraint method, weighted metric methods, value function methods, goal programming methods, and min-max optimization can only find a single solution from a single algorithm run, and some make assumptions about the shape and continuity of the Pareto frontier or require problem knowledge such as a set of suitable weights or target values (Deb, 2001; Coello Coello et al., 2007). Because of their advantages, many multi-objective evolutionary algorithms have been developed. Notable multi-objective evolutionary algorithms include PAES (Knowles and Corne, 1999), SPEA (Zitzler and Thiele, 1999), PESA (Corne et al., 2000), SPEA2 (Zitzler et al., 2001), NSGA-II (Deb et al., 2002), MOEA/D (Zhang and Li, 2007), SMS-MOEA (Beume et al., 2007), and NSGA-III (Deb and Jain, 2014).

To our knowledge, multi-objective evolutionary algorithms have not been employed in solving multi-objective optimization problems in plant breeding literature. Evolutionary algorithms have only been applied to classical multi-objective optimization techniques. For example, Akdemir and Sánchez (2016) used a hybrid genetic algorithm with a tabu search function to map Pareto frontiers using the weighted sum method. Gorjanc and Hickey (2018) used a modified differential evolution algorithm to solve for multiple ε-constraint problems and map a Pareto frontier for mating plans in their software package AlphaMate. Gorjanc et al. (2018) later used the differential evolution algorithm in AlphaMate to optimize cross selections in their simulations. Allier et al. (2019a) also used a differential evolution algorithm to solve a multi-objective mate selection problem which had been reduced to a single objective using the ε-constraint method. Finally, Butoto et al. (2022) and Cowling et al. (2023) both used evolutionary algorithms to solve for optimal contribution selection problems.

This paper presents Python Breeding Optimizer and Simulator (PyBrOpS), a Python package capable of performing numerical optimization for problems in breeding and simulating breeding pipelines. Several breeding simulation frameworks have been developed in years prior including AlphaSim (Faux et al., 2016), AlphaSimR (Gaynor et al., 2021), DeltaGen (Jahufer and Luo, 2018), ADAM-Plant (Liu et al., 2019), MoBPS (Pook et al., 2020), QuLinePlus (Hoyos-Villegas et al., 2019), and XSimVersion 2 (Chen et al., 2022). PyBrOpS is unique among these packages in that it is explicitly designed to simulate multiple traits and estimate Pareto frontiers for multiple breeding objectives.

## 2. Methods

### 2.1. Representation of a breeding program within PyBrOpS

Breeding programs are complex entities that can vary greatly based on a program’s resource availability, breeding goals, and the species being bred. To properly constitute or simulate any breeding program, it is necessary to develop a formal description of the breeding program that is comprehensive, unambiguous, and reproducible (Simianer et al., 2021). In PyBrOpS, breeding programs are represented using Operators and Protocols. An Operator represents a process in a breeding program that takes an entire breeding program state, transforms it, and returns a modified breeding program state (Figure 1A). Operators are meant to be broad in functionality to encompass the wide diversity of breeding programs. A Protocol represents a process in a breeding program that takes several inputs and returns a single output (Figure 1B). Protocols are meant to be narrow and specific in their functionality and used to construct the processes defined by Operators (Figure 2). Operators, once defined, can be assembled into a Universal Breeding Algorithm from which a breeding program can be simulated (Figure 2).

**Figure 1:**
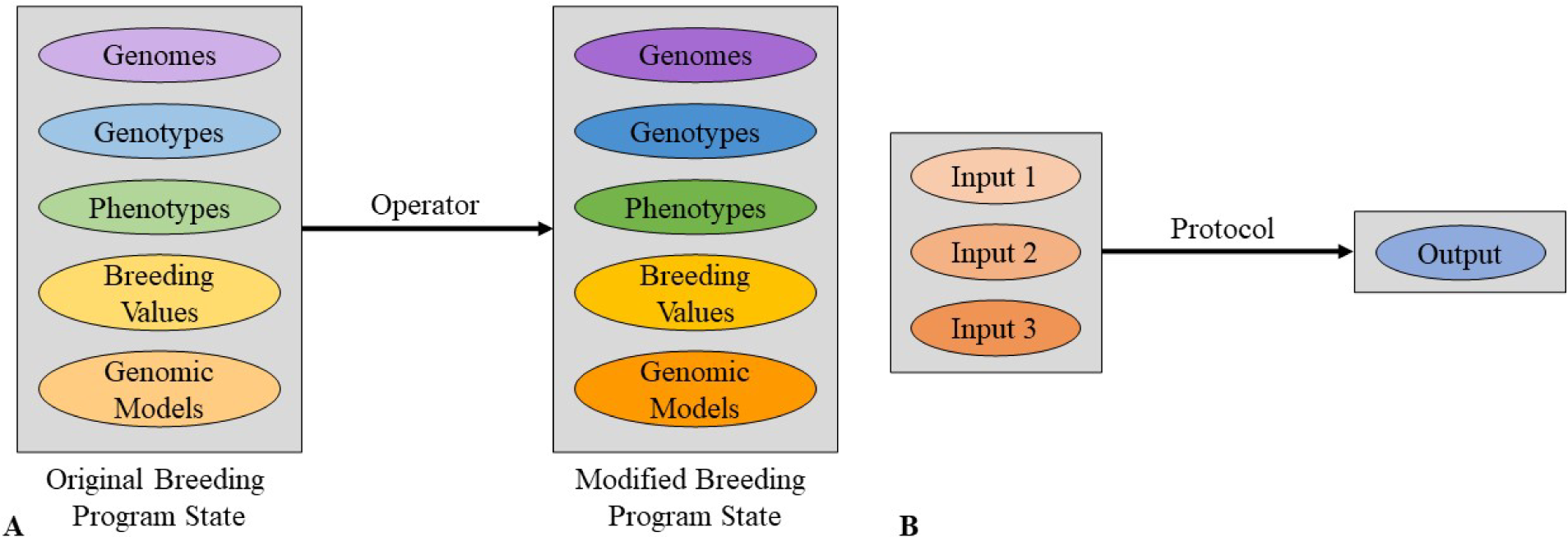
A) Breeding operators take entire breeding program states, which may consist of genomic, genotypic, phenotypic, breeding value, genomic model, and/or other types of data, and return a modified breeding program state. B) Breeding Protocols take data inputs and return a single output. The precise data types for each input and the output depend on the type of Breeding Protocol being used.

**Figure 2:**
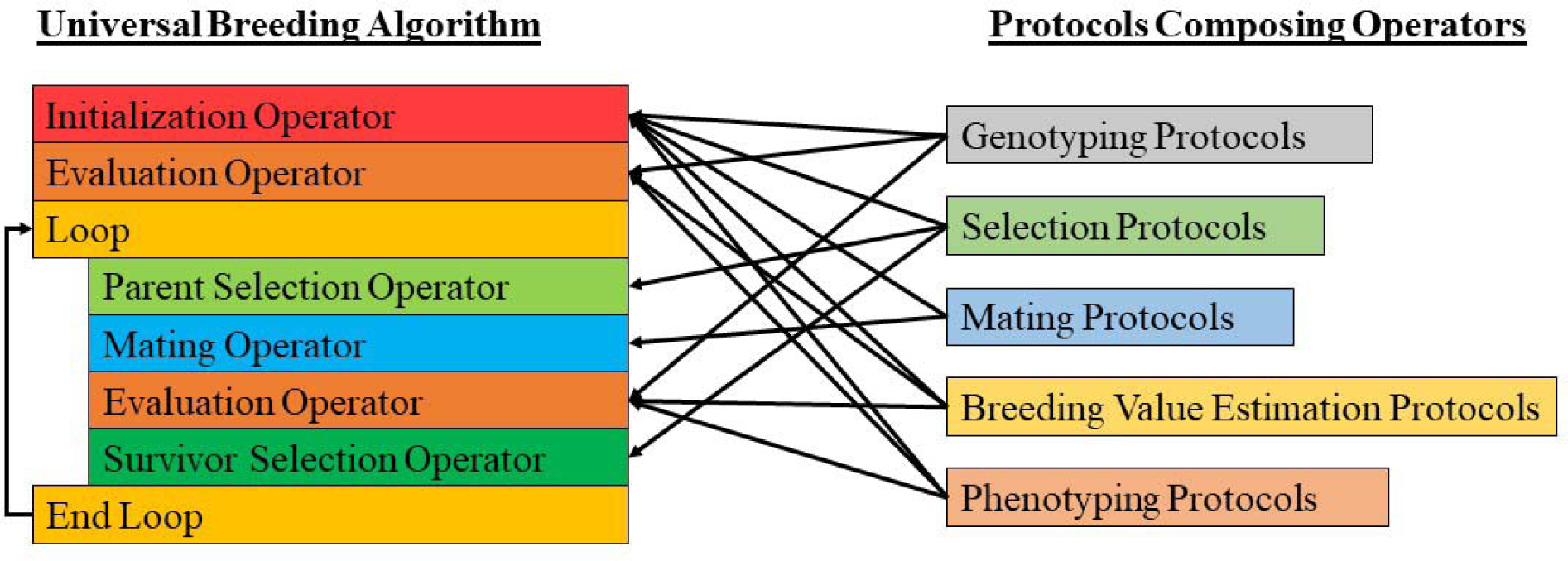
Left: Breeding Operators can be assembled into a Universal Breeding Algorithm. Outputs from each Operator are used as inputs for the next Operator to simulate a breeding program. Right: Breeding Protocols are modular and can be used to construct Operators and define their functionality.

In PyBrOpS, there are five defined Operators: Initialization, Parental Selection, Mating, Evaluation, and Survivor Selection (Figure 2). Briefly, each Operator’s purpose is described. The purpose of the Initialization Operator is to give a breeding program simulation a starting point from which to proceed. This may be as simple as loading genotypic and phenotypic data into PyBrOpS from a real breeding program or it may define a set of burn-in generations preceding a breeding program simulation as is common in many research simulations (Sonesson et al., 2012; Liu et al., 2015; Müller et al., 2018; Akdemir et al., 2019; Allier et al., 2019a). The Evaluation Operator of a breeding program performs all tasks related to the measurement of the desirability of individuals in the breeding program. This encompasses the genotyping of individuals, the phenotyping of individuals in field trials for one or more traits using experimental design, the development and deployment of genomic prediction models, and the modeling and estimation of breeding values. The purpose of the Parental Selection operator is to utilize information provided by the Evaluation Operator to select from a list of candidates a set of parents and a corresponding cross configuration mating those parents together to generate offspring. The purpose of the Mating Operator is to mate parents using the cross configuration selected by the Parental Selection Operator, simulate recombination, and generate progenies according to its instruction. Finally, the purpose of the Survivor Selection Operator is to use the information provided by the Evaluation Operator to select a set of individuals to serve as parental candidates for the next breeding cycle. This operator is like the Parent Selection Operator, but its main purpose is to quickly narrow down the set of breeding candidates to a manageable quantity and merge breeding subpopulations within the program. An example of a Survivor Selection Operator might perform within-family truncating selection followed by merging of selected family members into a single population.

PyBrOpS defines five Protocols: Breeding Value Estimation, Genotyping, Mating, Phenotyping, and Selection. Briefly, each Protocol’s purpose is described. The purpose of the Breeding Value Estimation Protocol is to provide routines for estimating breeding values from phenotypic and genotypic data. Mixed model analysis is nascent in Python, but users may utilize the statsmodels package (Seabold and Perktold, 2010) to fit linear mixed models and define custom Breeding Value Estimation Protocols. The purpose of the Genotyping Protocol is to take genomic information and simulate genotyping of individuals in a breeding program. The Mating Protocol module simulates recombination and mating. PyBrOpS has protocols implemented for self-crosses, two-way crosses, three-way crosses, four-way crosses, and doubled haploid generation from two-, three-, and four-way crosses. Random or uncontrolled mating, as may be prevalent in some species, can be simulated manually using implemented modules. The purpose of the Phenotyping Protocol is to simulate phenotypes resulting from field trials given genomic information. PyBrOpS implements a basic multi-environment protocol without genotype by environment effects. Finally, the purpose of the Selection Protocol is to provide methods for selecting individuals based on their genetic and/or phenotypic merits.

### 2.2. PyBrOpS implementation

PyBrOpS is implemented in pure Python and uses SOLID design principles in its architecture (Martin, 2018). SOLID design principles are a set of best practices that provide a framework for making software packages that are maintainable, comprehendible, modular, and extensible (Martin, 2018). Following SOLID principles, all objects in PyBrOpS implement an object interface – a contract defining standard object method members and behaviors. This means that for almost all modules in PyBrOpS, if the user desires to provide additional or novel functionality, the user need only implement a new class deriving from the interface. Objects implementing the same interface may be used interchangeably since routines depend on the interface abstraction and not any given object implementation.

### 2.3. PyBrOpS functionality

#### 2.3.1. Genome and genotype representation

PyBrOpS represents and stores genotypes and genomes in Genotype Matrix objects, which serve as wrappers around dense, 8-bit integer NumPy arrays (Oliphant, 2006). Genotype Matrix objects store marker variant information (e.g. single nucleotide polymorphism data) for individuals, along with metadata like the names of individuals, physical and genetic positions for each variant, and the ploidy level represented by the data. Genotypes are stored in two dimensional arrays, with dimensions representing individuals and marker variants, while genomes are stored in three dimensional arrays, with dimensions representing individuals, marker variants, and the specific chromosome phase or molecule along which the marker variant lies. PyBrOpS has the capability to load external genomic and genotypic data from VCF files using cyvcf2 (Pedersen and Quinlan, 2017) and create Genotype Matrices from raw NumPy arrays. Furthermore, genome and genotype data may be imported from or exported to a binary HDF5 format for convenient creation of breakpoints in a breeding simulation. If the user desires to implement a custom representation of a genome or genotype matrix, the user may implement a new class derived from a shared Genotype Matrix interface defining standard functionality for all Genotype Matrices.

#### 2.3.2. Genetic map and genetic map function representation

PyBrOpS represents genetic maps using Genetic Map objects, which serve as wrappers around NumPy arrays. Genetic Map objects store physical positions along with their corresponding genetic map positions in Morgans and support linear spline construction and interpolation of genetic positions from physical positions. PyBrOpS can load and save genetic map positions from and to CSV-like formats. If the user desires additional functionality, the user may create a new class that implements the Genetic Map interface defining standard functionalities for all Genetic Map classes.

Associated with genetic maps are genetic map functions. PyBrOpS implements two Genetic Map Function classes representing the Haldane (Haldane, 1919) and Kosambi (Kosambi, 1943) map functions. These two functions can be used to simulate recombination or calculate marker covariance caused by recombination (Allier et al., 2019b).

#### 2.3.3. Simulation of recombination

Recombination is simulated in a two-step process. In the first step, recombination probabilities between successive genomic markers are pre-computed using genetic map distances and a genetic map function and stored in a vector 𝐩_𝐱𝐨_. In the second step, a vector of uniform random numbers, 𝐮, is generated such that 𝑢_𝑖_∼Uniform(0,1) ∀𝑖. If the 𝑖th random number is less than the 𝑖th recombination probability, 𝑢_𝑖_ < (𝑝_𝑥𝑜_)_𝑖_, then a crossover occurs between the successive markers. Corresponding genome information is copied to progeny chromosomes.

#### 2.3.4. Genomic model representation

PyBrOpS represents genomic models using Genomic Model classes that provide model fitting and multi-trait prediction functionality. PyBrOpS implements two linear genomic models that follow Fisher’s model for quantitative genetics: a strictly additive linear genomic model, and an additive + dominance linear genomic model. The general mathematical representation of these models is as follows. Let 𝑛 be the number of individuals, 𝑡 be the number of traits, 𝑞 be the number of fixed effect predictors, and 𝑝 be the number of random effect predictors. Let 𝐘 be a 𝑛 × 𝑡 matrix of response variables, 𝐗 be a 𝑛 × 𝑞 matrix of fixed effect predictors, 𝐁 be a 𝑞 × 𝑡 matrix of fixed effect regression coefficients, 𝐙 be a 𝑛 × 𝑝 matrix of random effect predictors, and 𝐔 be a 𝑝 × 𝑡 matrix of random effect regression coefficients. Then the general mathematical representation of a linear genomic model is:

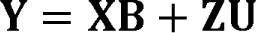

To model a strictly additive trait, we can interpret 𝐗 as a 𝑛 × 1 matrix of ones and 𝐁 as the intercept for each of 𝑡 traits. If we assume that elements in 𝐙 represent genotype calls coded as 0, 1, and 2, and elements in 𝐔 represent the effects of variants for each of 𝑡 traits, then the formula reduces to the familiar model for an additive trait:

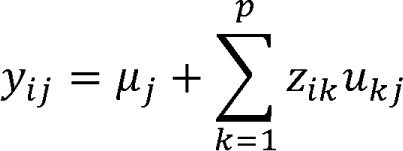

for the 𝑖th individual, for the 𝑗th trait.

To model an additive + dominance trait, we once again interpret 𝐗 as a 𝑛 × 1 matrix of ones, and 𝐁 as the intercept for each of 𝑡 traits. 𝐙 is interpreted as a block matrix composed of an 𝑛 × 𝑝_𝑎_ genotype block (𝐙_𝐚_), and an 𝑛 × 𝑝_𝑑_ dominance indicator block (𝐙_𝐝_):

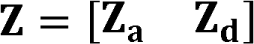

Elements in 𝐙_𝐚_ represent genotype calls as 0, 1, and 2. Elements in 𝐙_𝐝_ represent binary, indicator variables for whether an individual is heterozygous at a particular locus. Specifically, elements of 𝐙_𝐝_ are determined as (𝑧_𝑑_)_𝑖𝑘_ = 𝐼_((𝑧𝑎)𝑖𝑘=1)_((𝑧_𝑎_)_𝑖𝑘_). The 𝐔 matrix is interpreted as a block matrix containing 𝑝_𝑎_ × 𝑡 additive effects (𝐔_𝐚_) and 𝑝_𝑑_ × 𝑡 dominance effects (𝐔_𝐝_) for each of 𝑡 traits:

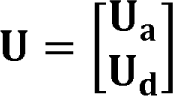

Putting all this together, then the formula reduces to the familiar model for an additive trait with dominance effects:

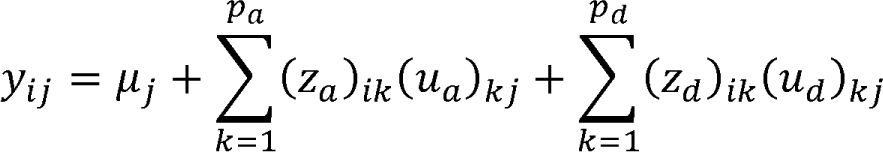

for the 𝑖th individual, for the 𝑗th trait.

Models may be imported from and exported to Pandas DataFrames, CSV-like formats, and HDF5 files, allowing users to use models fit with popular R packages like rrBLUP (Endelman, 2011) or BGLR (Pérez and de los Campos, 2014). Epistatic and nonlinear models are not implemented in PyBrOpS, but using the Genomic Model interfaces, one may implement custom epistatic or nonlinear models.

#### 2.3.5. Breeding value representation

PyBrOpS represents breeding values with Breeding Value Matrix objects. Breeding Value Matrix objects are wrappers around NumPy arrays and store relevant information such as names of the traits and individuals whose breeding values are stored in the matrix. PyBrOpS can load and save breeding values from Pandas DataFrames, CSV-like formats, and HDF5.

#### 2.3.6. Single- and multi-objective selection

PyBrOpS implements several Selection Protocols including conventional genomic selection (Meuwissen et al., 2001), conventional phenotypic selection, within family phenotypic selection, optimal contribution selection (Meuwissen, 1997), optimal haploid value selection (Daetwyler et al., 2015), optimal population value selection (Goiffon et al., 2017), expected maximum breeding value selection (Müller et al., 2018), weighted genomic selection (Jannink, 2010), and random selection. Selection Protocols provide four methods useful for making selections in physical breeding programs and in-silico breeding simulations.

The first method is the `problem` method. The `problem` method constructs a Selection Problem object from provided genomes, genotypes, phenotypes, breeding values, and/or a genomic prediction model.

Selection Problems define an optimization problem and encapsulate all required data and routines required to conduct optimizations for the defined problem. Selection Problems define the following optimization problem:

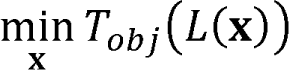

Such that:

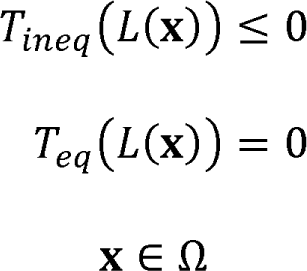

Here, 𝐱 is a decision vector that represents a selection decision in some selection decision space Ω. The meaning of selection decision depends on the context. If 𝐱 is a subset, it represents the subset of individuals or crosses chosen for selection. If 𝐱 is binary valued, it represents whether individuals or crosses are selected (1) or not (0). If 𝐱 is integer-valued, it represents an integer-valued contribution of an individual or cross to selection. Finally, if 𝐱 is real-valued, it represents a real-valued contribution of an individual or cross to selection. Next, 𝐿(𝐱) is a latent function that maps a selection decision to an 𝑙-dimensional latent vector: 𝐿(𝐱) ↦ ℝ^𝑙^. The latent function is unique to the Selection Problem and its dimension is determined by the Selection Protocol constructing the Selection Problem. The latent vector contains all objective and constraint evaluation information for a given selection strategy. Finally, 𝑇_𝑜𝑏𝑗_, 𝑇_𝑖𝑛𝑒𝑞_, and 𝑇_𝑒𝑞_ are transformation functions provided by the user to transform the latent vector into objectives, inequality constraints, and equality constraints, respectively. By providing different transformation functions, the user may convert various elements of the latent vector into constraints, or even new objectives. Latent vector transformation functions are transmitted from the Selection Protocol object to the Selection Problem object when the former constructs the latter. Selection Problem objects have methods to evaluate candidate solutions, allowing the user to manually evaluate the desirability of a proposed solution. All Selection Problem objects are also compatible with PyMOO (Blank and Deb, 2020) and can be solved with PyMOO infrastructure if desired.

The second method is the `sosolve` method. The `sosolve` method, given genomes, genotypes, phenotypes, breeding values, and/or a genomic prediction model, first creates a Selection Problem and then, assuming that the Selection Problem is single-objective in nature, solves it using a single-objective optimization algorithm. A generic genetic algorithm is the default optimization algorithm, but the user may utilize pre-built hillclimber algorithms or provide a custom single-objective optimization algorithm. Similarly, the third method, `mosolve`, creates a Selection Problem and solves the Selection Problem with a multi-objective optimization algorithm, assuming that the Selection Problem is multi-objective in nature. A generic NSGA-II (Deb et al., 2002) algorithm is the default multi-objective optimization algorithm, but the user may also utilize NSGA-III (Deb and Jain, 2014) or provide a custom multi-objective optimization algorithm to solve the problem. Both the `sosolve` and `mosolve` methods return a Selection Solution object, which encapsulates the solution or solutions to the optimization problem.

The final method is the `select` method. Given genomes, genotypes, phenotypes, breeding values, and/or a genomic prediction model, the `select` method constructs a Selection Problem, and solves the problem using an appropriate single- or multi-objective problem. If the problem is single-objective, it constructs and returns a Selection Configuration object from the singular solution obtained from the optimization. If the problem is multi-objective, then the solution is a set of non-dominated solutions.

Since selection necessitates that a single decision is made, solutions in the non-dominated set are transformed by a non-dominated set transformation function 𝑇_𝑁𝐷_. This function accepts objective coordinates for the whole non-dominated set and assigns a single score to each solution: 𝑇_𝑁𝐷_(ℝ^𝑛×𝑑^) ↦ ℝ^𝑛^. In some applications, this may be a function that calculates the distance between a preference weight vector and a non-dominated point. The solution with the best score is used to construct a Selection Configuration object. Selecting points from a non-dominated set of solutions may be useful in situations where the Pareto frontier is non-convex. In situations where the Pareto frontier is non-convex, simple linear weighting methods will fail to find points in non-convex portions of the Pareto frontier (Deb, 2001). In both single- and multi-objective selection scenarios, the returned Selection Configuration object encapsulates all information necessary to identify the individuals or crosses that should be made. This information can be used to drive simulations or inform the breeder.

## 3. Results

To demonstrate the capabilities of PyBrOpS, three use-case scenarios are exhibited. In scenario 1, a Pareto frontier is mapped for Optimal Contribution Selection (OCS; Meuwissen, 1997) with two simulated quantitative traits. In scenario 2, a Pareto frontier for OCS with two real-world yield traits in wheat is mapped from which a single solution is selected. In scenario 3, Pareto frontiers are mapped and subsequently used to make selections for two simulated quantitative traits using Conventional Genomic Selection (Meuwissen et al., 2001).

### 3.1. Example Scenario 1: Mapping a Pareto frontier for Optimal Contribution Selection with two simulated quantitative traits

In the first use-case scenario, 2000 markers were randomly selected with minor allele frequencies greater than 0.2 from the Wisconsin Maize Diversity Panel 942 (Hirsch et al., 2014; Mazaheri et al., 2019) to serve as empirical marker sources. The US NAM maize linkage map (McMullen et al., 2009) was used to linearly interpolate genetic map positions for each of the 2000 selected markers. For each marker, strictly additive effects were assigned for two quantitative traits following a multivariate normal distribution with negative covariance to simulate two traits that are competing in nature:

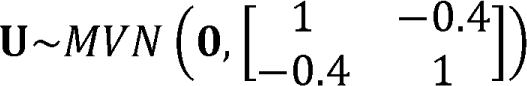

The intercepts for quantitative traits 1 and 2 were set to 10 and 25, respectively.

A multi-objective optimization of a subset variant of OCS (Meuwissen, 1997) was conducted and can be summarized as follows:

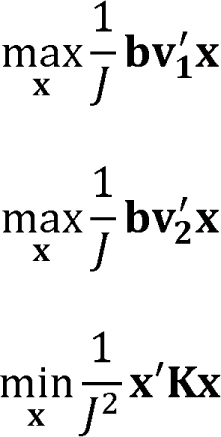

Such that:

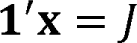

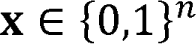

Here, 𝐱 represents a subset of 𝐽 = 20 individuals selected from the set of candidates in the diversity panel. 𝐛𝐯_𝟏_ and 𝐛𝐯_𝟐_ represent the true breeding values of panel members for quantitative traits 1 and 2, respectively, using the assigned genomic model. 𝐊 represents a matrix of identity-by-state kinship relationships between individuals, calculated using methods described by Allier et al., 2019a. The first two objectives can be interpreted as maximizing the mean breeding value for quantitative traits 1 and 2 in the selected subset. The third objective can be interpreted as minimizing the average kinship between individuals in the selected subset.

An NSGA-III (Deb and Jain, 2014) algorithm with a population size of 100 and 91 reference points was evolved for 1500 generations to approximate the Pareto frontier for the problem formulated above and identify a set of relatively evenly distributed non-dominated points for each of the three objectives (Figure 3). While appearing to be smooth and convex from a distance, the estimated Pareto frontier contains regions that are locally non-convex. These local non-convexities are likely due to the discrete nature of the optimization problem but could also be attributable to premature convergence to the Pareto frontier.

**Figure 3:**
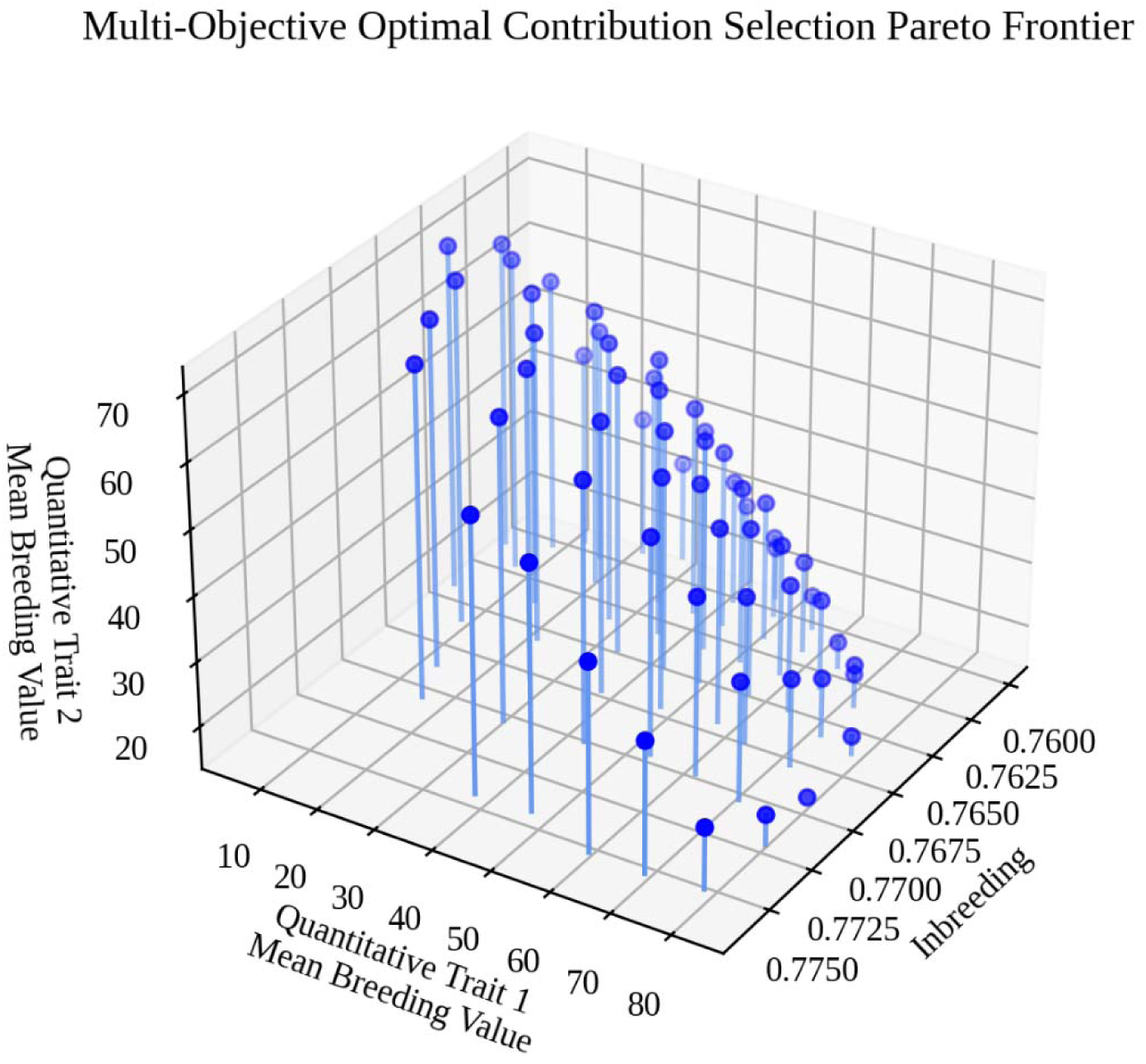
A three-dimensional Pareto frontier for the OCS problem in Scenario 1, visualized using Matplotlib (Hunter, 2007) in Python. The x-axis represents mean inbreeding of the selected subset, the y-axis represents the mean breeding value (genetic gain) of the selected subset for quantitative trait 1, and the z-axis represents mean breeding value (genetic gain) of the selected subset for quantitative trait 2.

### 3.2. Example Scenario 2: Mapping a Pareto frontier for Optimal Contribution Selection with two real-world yield traits in wheat

In the second use-case scenario, a multi-objective optimization for OCS with two traits was performed using a historical CIMMYT wheat dataset from the BGLR package (Pérez and de los Campos, 2014) as an empirical source of genomic markers and phenotypic data. Specifically, the dataset contained 1279 markers and yield data in four locations for 599 unique individuals. We took these data, fit an RR-BLUP model for two of the traits using the rrBLUP package in R (Endelman, 2011), and exported the results as a set of files. Using the exported files, we loaded marker and model information into PyBrOpS, calculated genomic estimated breeding values (GEBVs) for the dataset, and performed a multi-objective optimization using a problem formulation and multi-objective optimization strategy identical to that which is described in Scenario 1. After performing the optimization (Figure 4), we selected a point corresponding to a preference of 0.2 for inbreeding, 0.4 for the first trait, and 0.4 for the second trait from the estimated Pareto frontier using a pseudoweight method. Briefly, the objective values for points in the estimated Pareto frontier were scaled to the range [0,1]. For each point, the Euclidean distance between the point’s normalized values and a [0.2, 0.4, 0.4]^′^ preference vector originating from the origin was calculated. The point with the smallest Euclidean distance was selected as the preferred selection solution for the problem (Figure 4; highlighted in red).

**Figure 4:**
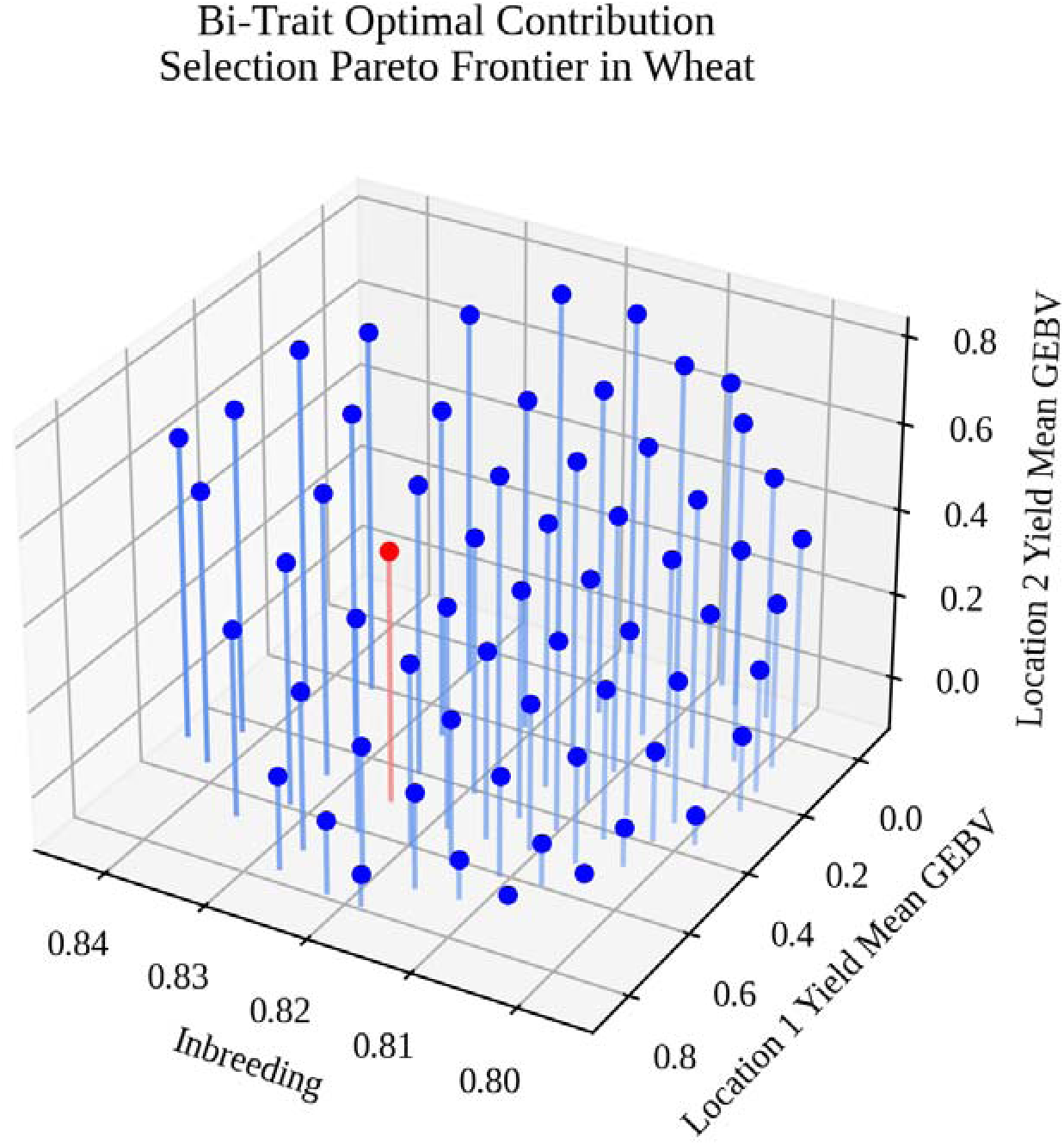
A three-dimensional Pareto frontier for the bi-trait OCS problem with the wheat dataset. The x-axis represents mean inbreeding of the selected subset, the y-axis represents the mean GEBV of the selected subset for yield at location 1, and the z-axis represents mean GEBV of the selected subset for yield at location 2. Highlighted in red is the selection decision corresponding to a pseudoweight preference of 0.2 for inbreeding, 0.4 for yield at location 1, and 0.4 for yield at location 2.

### 3.3. Example Scenario 3: Multi-objective breeding program simulation for two simulated quantitative traits

In the third use-case scenario, we used the same base simulation described in Scenario 1 except 1000 markers were sampled to simulate quantitative traits instead of 2000. Breeding program simulations were performed following a protocol like what is described in Allier et al. (2019a). We wrote a Python script to perform a burn-in in PyBrOpS. Briefly, 40 unique, randomly selected genotypes were randomly intermated for 10 generations to create a founder population for the breeding simulation. Narrow sense heritabilities of 0.4 and 0.6 for a single simulated environment were assigned to quantitative traits 1 and 2, respectively, by calculating the trait genetic variances in the base population and backsolving for the error variances assuming no genotype by environmental interactions. Environmental variances were assumed to be constant throughout simulations. As a result, reductions in genetic diversity resulting from selection reduced the narrow sense heritability as the simulations progressed. The simulation was initiated with 10 generations of phenotypic selection for burn-in to mimic linkage disequilibrium patterns that resemble those found in real breeding programs. In each iteration of the burn-in stage, progenies were evaluated in 4 simulated locations assuming no genotype by environment interactions. Phenotypic values were simulated as 𝑦_𝑖𝑗𝑘_ = 𝑔_𝑖𝑘_ + 𝑙_𝑗𝑘_ + 𝜀_𝑖𝑗𝑘_ where 𝑦_𝑖𝑗𝑘_ is the phenotype of the 𝑖th individual in the 𝑗th location for the 𝑘th trait, 𝑔_𝑖𝑘_ is the genotypic value of the 𝑖th individual for the 𝑘th trait calculated according to the established model, 𝑙_𝑗𝑘_ is the location mean in the 𝑗th location for the 𝑘th trait, and 𝜀_𝑖𝑗𝑘_ is the phenotypic error of the 𝑖th individual in the 𝑗th location for the 𝑘th trait. Location means were set as zero for all locations and traits. Phenotypic errors were sampled as 𝜀_𝑖𝑗𝑘_∼𝑁(0, 𝜎^2^) where 𝜎^2^ is the error variance for the 𝑘th trait established from the founder population. Breeding values for individuals were calculated by taking the mean across locations. The top 5% (4 individuals) from each family, according to the sum of estimated breeding values, were selected to serve as parental candidates for the next generation. After narrowing the list of parental candidates down to a manageable level, the top 40 individuals according to the sum of estimated breeding values were selected as parents and randomly intermated in 20 biparental crosses to produce 80 progenies per cross. Parental candidates from the previous three generations were considered in the selection of founders for the next generation.

After the burn-in, the true genomic model was used to make selections using conventional genomic selection. The main simulation proceeded in a manner identical to the burn-in except for the parental selection step. Instead, parental selections were made by approximating the Pareto frontier using NSGA-II (Deb et al., 2002), scaling trait value ranges of the identified Pareto optimal set to the range [0,1], and selecting the solution closest to a weight of 0.5 (equal preference) for both scaled traits. The selected solution contained information on which 40 parents were to be selected as founders for the next generation. The simulation lasted 10 generations. Throughout the simulation, the Pareto frontier at each selection point was saved and mean population trait breeding values were recorded. The Pareto frontiers and mean population breeding values throughout time were plotted using Matplotlib (Hunter, 2007) and are visualized in Figures 5A and 5B, respectively.

**Figure 5:**
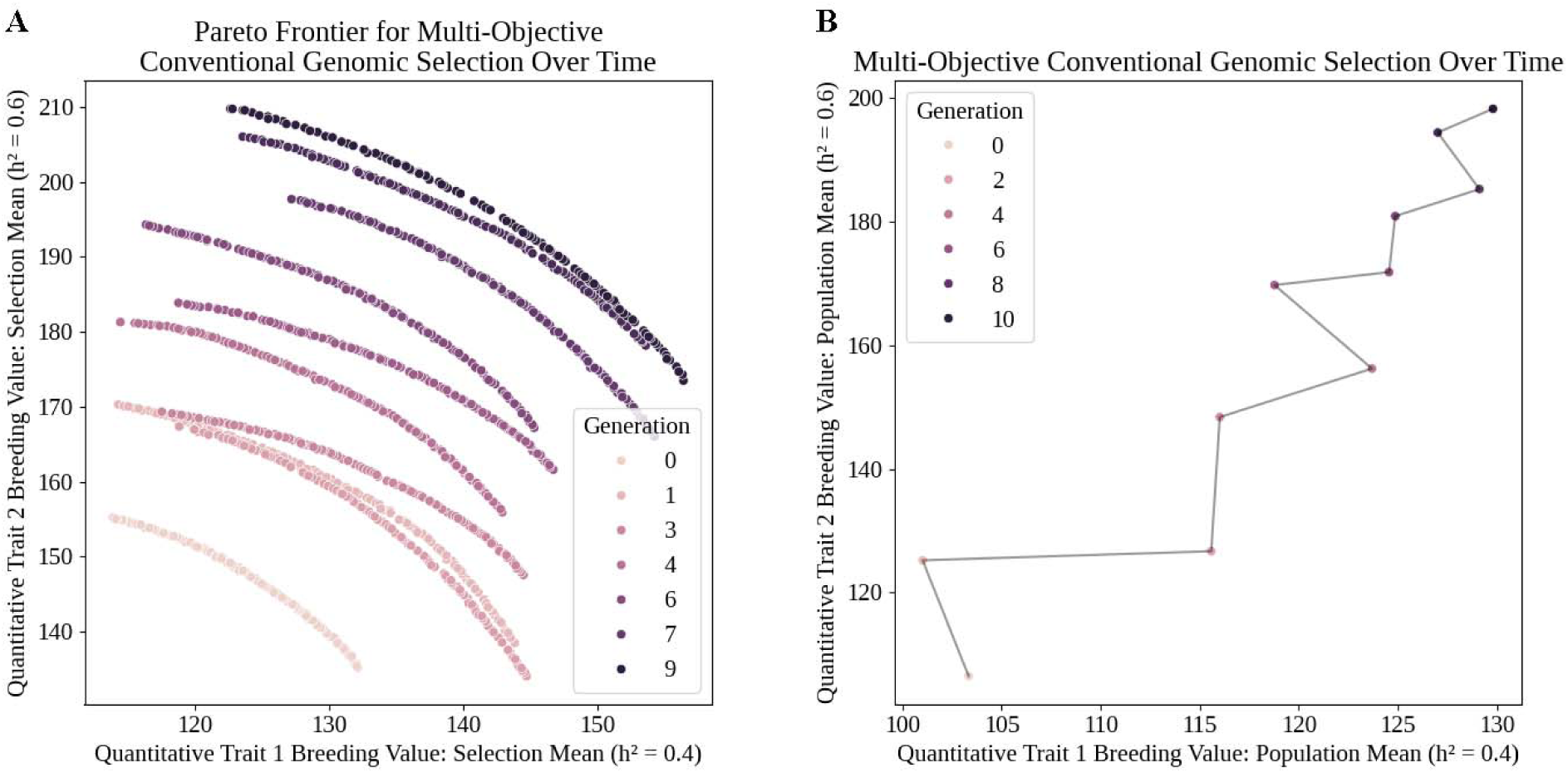
A) Pareto frontiers mapped by NSGA-II and B) population mean breeding values at each generation in Scenario 2 simulations. A) The Pareto frontier at Generation 0 represents the initial population selection possibilities, while the frontier at Generation 9 represents the selection possibilities before the final mating round in the simulation. Individual points along each Pareto frontier represent potential subsets of individuals a breeder could select. B) Mean trait breeding values for the breeding population in Scenario 2 simulations.

## 4. Conclusion

PyBrOpS is a Python package for simulating breeding programs and performing multi-objective optimizations. PyBrOpS has the capability to simulate multiple quantitative traits, simulate the evolution of entire breeding programs, and map breeding frontier possibilities for these traits. Stochastic simulations serve as a valuable tool in a breeding program and can be used to optimize a pipeline for genetic gain, genetic diversity, economic value, or other program objectives. Additionally, the native ability of PyBrOpS to map breeding frontiers for multiple traits is of particular interest to breeders, decision-makers, and other stakeholders. Mapping breeding frontiers facilitates the visualization and understanding of tradeoffs and can assist in decision making.

Due to the high degree of modularity built into PyBrOpS’s architecture, new features are easy to implement. To provide additional functionality, the user may implement an interface relevant to the feature desired to be added. Future versions of PyBrOpS will include additional selection protocols and offer genomic prediction routines native to Python.

## 5. Web resources

PyBrOpS is open source and can be found on GitHub at https://github.com/rzshrote/pybrops/. Complete documentation, examples from this paper, and examples for other multi-objective optimization and simulation scenarios can be found on the associated GitHub Pages website at https://rzshrote.github.io/pybrops/. Feature requests, bug reports, and feature submissions may be made at the aforementioned GitHub repository link.

## Acknowledgements

This work is supported in part by the National Science Foundation Research Traineeship Program (DGE 1828149) and the MSU Plant Science Fellowship awarded to RZS. RZS would like to acknowledge and thank his father Curtis Shrote for recommending resources and sharing his wisdom about software architecture.

